# Necessity and Impact of Specialization of Large Foundation Model for Medical Segmentation Tasks

**DOI:** 10.1101/2024.06.02.597036

**Authors:** Eric Nguyen, Hengjie Liu, Dan Ruan

## Abstract

**Background:** Large foundation models, such as the Segment Anything Model (SAM), have shown remarkable performance in image segmentation tasks. However, the optimal approach to achieve true utility of these models for domain-specific applications, such as medical image segmentation, remains an open question. Recent studies have released a medical version of the foundation model MedSAM by training on vast medical data, who promised SOTA medical segmentation. Independent community inspection and dissection is needed.

**Purpose:** This study assesses the performance of off-the-shelf medical foundation model MedSAM for the segmentation of anatomical structures in pelvic MR images. We also evaluate the dependency on prompting scheme and demonstrate the gain of further specialized fine-tuning.

**Methods:** MedSAM and its lightweight version LiteMedSAM were evaluated out-of-the-box on a public MR dataset consisting of 589 pelvic images split 80:20 for training and testing. An nnU-Net model was trained from scratch to serve as a benchmark and to provide bounding box prompts for MedSAM. MedSAM was evaluated using different quality bounding boxes, those derived from ground truth labels, those derived from nnU-Net, and those derived from the former two but with 5-pixel isometric expansion. Lastly, LiteMedSAM was refined on the training set and reevaluated on this task.

**Results:** Out-of-the-box MedSAM and LiteMedSAM both performed poorly across the structure set, especially for disjoint or non-convex structures. Varying prompt with different bounding box inputs had minimal effect. The mean Dice score and mean Hausdorff distances (in mm) for obturator internus using MedSAM and LiteMedSAM were {0.251 ± 0.110, 0.101 ± 0.079} and {34.142 ± 5.196, 33.688 ± 5.306}, respectively. Fine-tuning of LiteMedSAM led to significant performance gain, improving Dice score and Hausdorff distance for the obturator internus to 0.864 ± 0.123 and 5.022 ± 10.684, on par with nnU-Net with no significant difference in evaluation of most structures. All segmentation structures all benefited significantly from specialized refinement, at varying improvement margin.

**Conclusion:** Our study alludes to the potential of deep learning models like MedSAM and lite MedSAM for medical segmentation but also highlight the need for specialized refinement and adjudication: it is quite likely that off-the-shelf use of such large foundation models may be suboptimal, and specialized fine-tuning can significantly enhance segmentation accuracy.

## 1. Introduction

Foundation models are general models trained on extremely large and diverse datasets that make up a wide range of categories which enable the model to be multi-purpose.^1^ Segment Anything (SAM) is a generalist foundation model developed by Meta for image segmentation. It is trained on over 11 million natural images with 1 billion masks and has shown remarkable zero-shot segmentation performance on a diverse range of image segmentation tasks.^2,3^ SAM utilizes a variety of prompts to infer segmentation intention. Studies have shown that SAM may be challenged by low image contrast and amorphous target structures in medical image segmentation, resulting in much poorer and unstable performance compared to natural image applications.^4–7^ Multiple methods have been devised to bridge this gap such as the incorporation of Low Rank Adaptation (LoRA) into SAM’s image encoder, a technique that has previously shown great potential for other large pre-trained models.^4,8,9^ SAM-Adapter was developed primarily for improved detection of shadows and camouflaged objects in natural images, but has also been shown to improve performance of polyp segmentation for medical images.^10,11^ Another study proposed a 3D adaptation of SAM, originally limited to 2D images, to better facilitate medical images.^5^ Ma *et al*. preserves the architecture of the original SAM and instead provides a new refined foundation model, MedSAM, by fine-tuning SAM on an unprecedented dataset with over a million annotated medical images.^12^

MedSAM was trained predominantly on MR and CT images, but the training set also spans endoscopy, ultrasound, x-ray, pathology, fundus, dermoscopy, and OCT. The authors report that MedSAM shows significant improvement over SAM for medical applications and can perform on par with modality-specific state-of-the-art models. In this study, we investigate and assess the clinical feasibility of MR-based pelvic segmentation using various variants in the MedSAM family. Specifically, we evaluate the performance of off-the-shelf MedSAM, its dependency and sensitivity on the prompt input, and impact of fine tuning. Comparison was performed against a special-purpose in-house trained nnU-Net benchmark.

## 2. Material and Methods

### 2.1 Dataset description

We used a dataset curated by Li et al. consisting of 589 T2w MRI images acquired from seven studies (INDEX, the SmartTarget Biopsy Trial, PICTURE, TCIA Prostate3T, Promise12, TCIA ProstateDx and the Prostate MR Image Database).^13^ All the studies are represented equally in the total curated dataset. The collection contains images from multiple institutions that were done using scanners with either 1.5T or 3.0T field strengths and from two different manufacturers. The images had an in-plane resolution ranging from 0.3 mm to 1.0 mm and a slice thickness ranging from 1.8 mm to 5.4 mm and had varying field-of-views. All images distributed by the curated dataset had dimensions of 180×180×N pixels. Manually annotated segmentations were also provided for each of the images. Eight anatomical structures were labelled, including bladder, bone, obturator internus (oi), transition zone, central gland, rectum, seminal vesicle (sv), and neurovascular bundle (nvb).

The current SAM and MedSAM can only cope with 2D segmentation, so slices in each 3D volume are processed and segmented independently and then recompiled back into the volume space for performance evaluation.

### 2.2 Background on SAM and MedSAM

MedSAM consists of a vision transformer (ViT)-based encoder for feature extraction, a prompt encoder that accepts user-provided bounding box inputs, and a lightweight mask decoder. MedSAM uses image inputs of size 1024x1024 while LiteMedSAM uses image inputs of size 256×256. Our images were resampled to meet the respective dimension criteria.

MedSAM was pre-trained using a loss function defined as the unweighted sum of Dice loss and binary cross entropy (BCE) loss. Meanwhile, the loss function used for the pre-training of LiteMedSAM also includes intersection over union (IoU) loss into this summation. The loss function used by LiteMedSAM is shown below. Let N be the total number of voxels in an image and let g_i_ and s_i_ be the ith voxel of the ground truth and predicted segmentations, respectively.

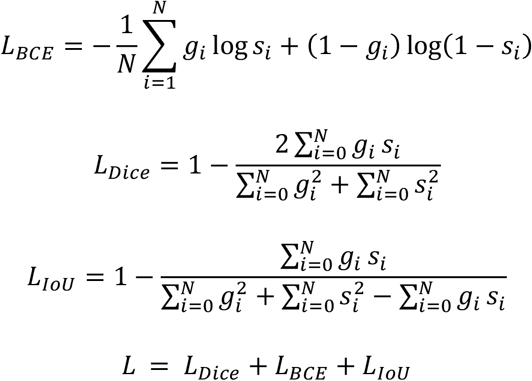

We examined the performance of the pre-trained MedSAM and a pre-trained LiteMedSAM on the testing data and task described in Section II.C.

### 2.3 Prompting Variations

MedSAM models take bounding box input as prompts. We have conducted investigation on four different options of bounding box prompts: oracle bounding boxes derived from the ground truth mask without and with 5-pixel isometric expansion, and bounding boxes derived from nnU-Net segmentation mask without and with 5-pixel isometric expansion. The prompts derived from ground truth labels are used to reflect the best-case prompting while those derived from nnU-Net labels simulate average-case prompting.

### 2.4 Refinement of Lite MedSAM with specialized data and task

For fine-tuning of LiteMedSAM, the original 589 3D MR images were randomly split at an 80:20 ratio for training and hold-off testing in this study. The training group was again randomly split at an 80:20 ratio for training and validation. In total, 18,350 2D image-mask pairs were used to train the LiteMedSAM model. Data augmentation was utilized during training which consisted of random left-right or up-down flips. During training, bounding boxes were also generated from the ground-truth masks and used as an additional model input. The bounding box inputs had 5-pixel random shift. The network was optimized using Adam optimizer (β_1_=0.9, β_2_=0.999) with a learning rate of 5e-5 and a weight decay of 0.01. The same loss function used for training of LiteMedSAM was again used here for fine-tuning.

For consistency with the preprocessing protocol described in the MedSAM paper, the following two exclusion criteria were applied on the training dataset. To improve dataset quality, tiny objects defined by a 100-pixel threshold were removed from slices as the challenge would be detection rather than segmentation. In addition, the intensity levels of all images were clipped between the 0.5th and 99.5th percentiles, and min-max normalization was applied.

Training was performed on a single GPU (NVIDIA GeForce RTX3080, 10G memory) using resampled image inputs of size 256×256 to accommodate the memory limit. The model typically converges in about 300 epochs with a batch size of 4. The checkpoint with the best validation loss was chosen for the final model.

### 2.5 Benchmark nnU-Net model

nnU-Net was trained from scratch and using the same dataset and dataset splitting as with LiteMedSAM fine-tuning. nnU-Net offers both 2D and 3D U-Net configurations for training on new datasets. We opted to use the “3D Full res” configuration which uses the native image resolution (180×180xN). The network was trained over 1000 epochs and the checkpoint with the best validation loss was chosen. This U-Net model serves two purposes: a benchmark for overall performance comparison, and a means to generate bounding box prompts for MedSAM. We opted to use nnU-Net because it has been shown to perform well on a large variety of segmentation tasks.^14–16^

### 2.6 Evaluation Metrics

Dice scores and Hausdorff distances were calculated for each anatomical structure segmented for all patients using each of the models. Comparisons between each of the models were performed using a two-sided paired t-test, on statistics generated from the testing of 117 volumetric images. A significance level of 0.05 was used.

## 3. Results

### 3.1 Suboptimal performance for off-the-shelf SAM models

Regardless of the intrinsic segmentation challenge level based on the scale and morphology of the structure or intensity context, MedSAM or LiteMedSAM is notably inferior compared to the specialized nnU-Net performance, as shown in *Figure 1*. The benchmark nnU-Net showed statistically significant superiority in Dice score for all structures (p < 0.0001) and in Hausdorff distance for all structures (p < 0.05) except seminal vesicle in which no significance difference was noted. Specifically, for structures with well-defined and reasonably convex shapes, as in bladder, rectum, and intact prostate, MedSAM resulted in dice scores of 0.808, 0.846, and 0.833, respectively while LiteMedSAM achieved comparable dice scores of 0.783, 0.787, and 0.818, respectively. That’s 10-20% lower than mean dice scores of 0.958, 0.924, and 0.918 from the benchmark nnU-Net. In addition, both MedSAM and LiteMedSAM experienced great difficulty segmenting disjoint, bilateral structures including femur, neurovascular bundle, and obturator internus. For these structures, both models yielded mean Dice scores lower than 0.6. The specific values are reported in columns 1, 5, and 7 in *Table 1*. Illustrative examples are shown in *Figure 2*.

**Table 1:** Mean (and standard deviation) Dice Scores and Hausdorff Distances for different models. Best value for each structure is bolded.

| Dice Scores |  |  |  |  |  |  |  |
| --- | --- | --- | --- | --- | --- | --- | --- |
| Model (bbox option) | MedSAM (oracle) | MedSAM (Unet) | MedSAM (oracle expanded) | MedSAM (Unet expanded) | LiteMedSAM (oracle) | LiteMedSAM – refined (oracle) | nnU-Net (none) |
| Dice Scores |  |  |  |  |  |  |  |
| bladder | 0.81 ± 0.24 | 0.80 ± 0.24 | 0.810 ± 0.24 | 0.80 ± 0.24 | 0.78 ± 0.25 | 0.92 ± 0.17 | **0.96 ± 0.06** |
| bone | 0.58 ± 0.19 | 0.57 ± 0.19 | 0.59 ± 0.18 | 0.58 ± 0.19 | 0.51 ± 0.19 | 0.92 ± 0.14 | **0.95 ± 0.05** |
| central gland | 0.84 ± 0.10 | 0.80 ± 0.09 | 0.87 ± 0.09 | 0.83 ± 0.09 | 0.80 ± 0.12 | **0.91 ± 0.08** | 0.90 ± 0.04 |
| intact prostate | 0.83 ± 0.09 | 0.81 ± 0.08 | 0.84 ± 0.09 | 0.82 ± 0.08 | 0.82 ± 0.09 | 0.90 ± 0.08 | **0.92 ± 0.04** |
| NVB | 0.29 ± 0.10 | 0.25 ± 0.10 | 0.33 ± 0.10 | 0.28 ± 0.10 | 0.22 ± 0.12 | **0.72 ± 0.13** | 0.71 ± 0.15 |
| OI | 0.25 ± 0.11 | 0.24 ± 0.11 | 0.29 ± 0.11 | 0.28 ± 0.11 | 0.10 ± 0.08 | 0.86 ± 0.12 | **0.89 ± 0.06** |
| rectum | 0.85 ± 0.12 | 0.83 ± 0.11 | 0.86 ± 0.11 | 0.84 ± 0.11 | 0.79 ± 0.12 | 0.91 ± 0.11 | **0.92 ± 0.05** |
| SV | 0.72 ± 0.16 | 0.64 ± 0.17 | 0.74 ± 0.16 | 0.65 ± 0.18 | 0.69 ± 0.16 | **0.81 ± 0.18** | 0.79 ± 0.12 |
| Hausdorff Distances |  |  |  |  |  |  |  |
| bladder | 6.63 ± 6.05 | 6.68 ± 5.50 | 6.55 ± 6.17 | 6.55 ± 5.54 | 7.53 ± 6.44 | 2.76 ± 4.70 | **1.97 ± 2.11** |
| bone | 56.15 ± 13.87 | 57.34 ± 13.13 | 55.80 ± 14.09 | 56.98 ± 13.34 | 57.37 ± 12.52 | 3.97 ± 14.11 | **2.77 ± 11.53** |
| central gland | 4.12 ± 3.20 | 4.53 ± 2.51 | 3.67 ± 3.27 | 4.12 ± 2.52 | 4.74 ± 2.94 | **2.64 ± 2.78** | 3.17 ± 1.67 |
| intact prostate | 6.82 ± 4.12 | 6.51 ± 3.02 | 6.83 ± 4.11 | 6.34 ± 2.35 | 6.75 ± 3.88 | 5.57 ± 4.58 | **3.49 ± 2.85** |
| NVB | 13.55 ± 5.35 | 13.43 ± 5.73 | 13.56 ± 5.34 | 13.39 ± 5.60 | 13.52 ± 5.47 | 5.72 ± 6.44 | **5.49 ± 6.30** |
| OI | 34.14 ± 5.20 | 33.66 ± 4.30 | 34.01 ± 5.33 | 33.50 ± 4.41 | 33.69 ± 5.31 | 5.02 ± 10.68 | **4.98 ± 5.13** |
| rectum | 8.39 ± 12.47 | 8.25 ± 8.14 | 8.26 ± 12.62 | 8.21 ± 8.20 | 9.48 ± 11.99 | 6.03 ± 12.56 | **5.10 ± 7.11** |
| SV | 6.40 ± 4.31 | 8.23 ± 6.50 | 6.30 ± 4.39 | 8.17 ± 6.65 | 5.52 ± 3.42 | **3.39 ± 3.19** | 4.95 ± 4.96 |

**Figure 1:**
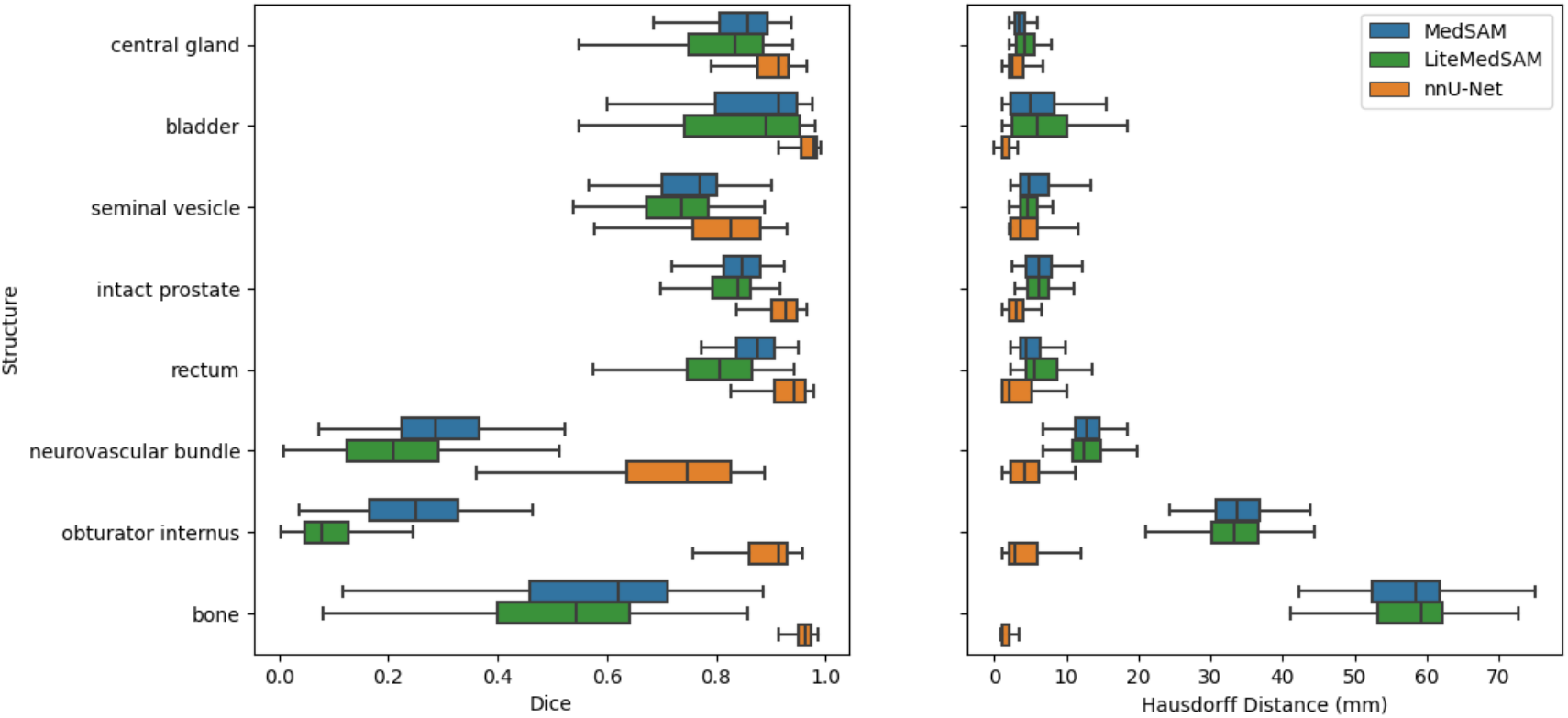
Comparison of Dice scores and Hausdorff distances among MedSAM (blue), LiteMedSAM (green), and nnU-Net (orange). While MedSAM and LiteMedSAM showcase comparable performance, nnU-Net is significantly superior.

**Figure 2:**
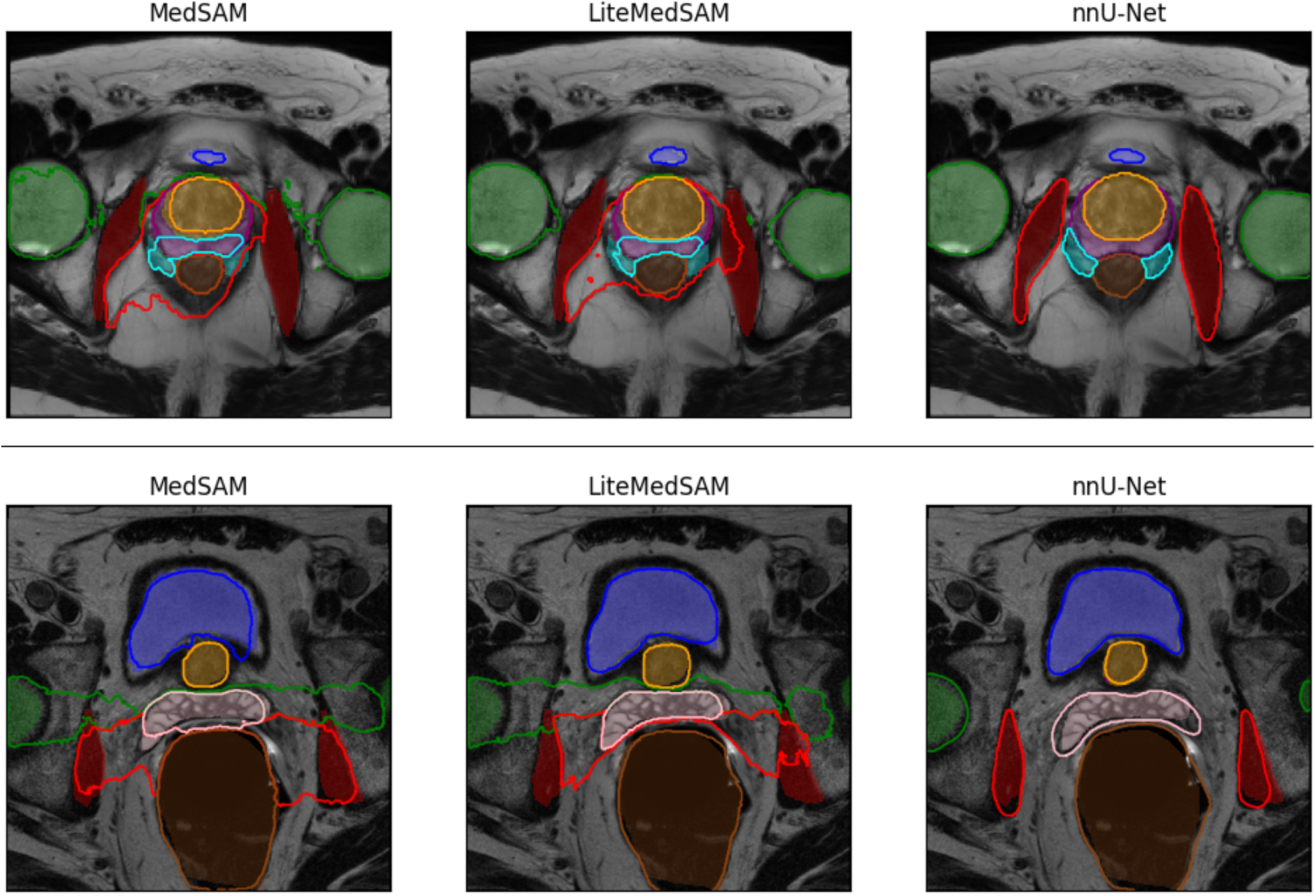
Examples of off-the-shelf MedSAM and LiteMedSAM segmentations compared to nnU-Net or two different patients. Shading indicates ground-truth. Structures shown: femur (green), bladder (blue), central gland (orange), intact prostate (purple), obturator internus (red), seminal vesicle (pink), neurovascular bundle (cyan), rectum (brown)

### 3.2 Impact of bounding box prompt

MedSAM models currently only offer stable support for prompts in the form bounding box. We evaluated the stability of MedSAM segmentations when provided with different bounding box prompts. To simulate best-case prompting, we provided MedSAM with oracle bounding boxes derived from the ground truth labels. Meanwhile, bounding boxes generated from nnU-Net was used to approximate average-case prompting. As shown in *Figure 3*, when provided bounding boxes generated by nnU-Net labels, MedSAM inferences saw a statistically significant decrease in mean Dice score compared to MedSAM inferences using ground-truth bounding boxes for prostate central gland (0.808 vs 0.800), intact prostate (0.836 vs 0.801), seminal vesicle (0.718 vs 0.637), and neurovascular bundle (0.294 vs 0.249). No significant difference in Hausdorff distances were observed between the two tests, except for seminal vesicle (6.34mm vs 8.23mm).

**Figure 3:**
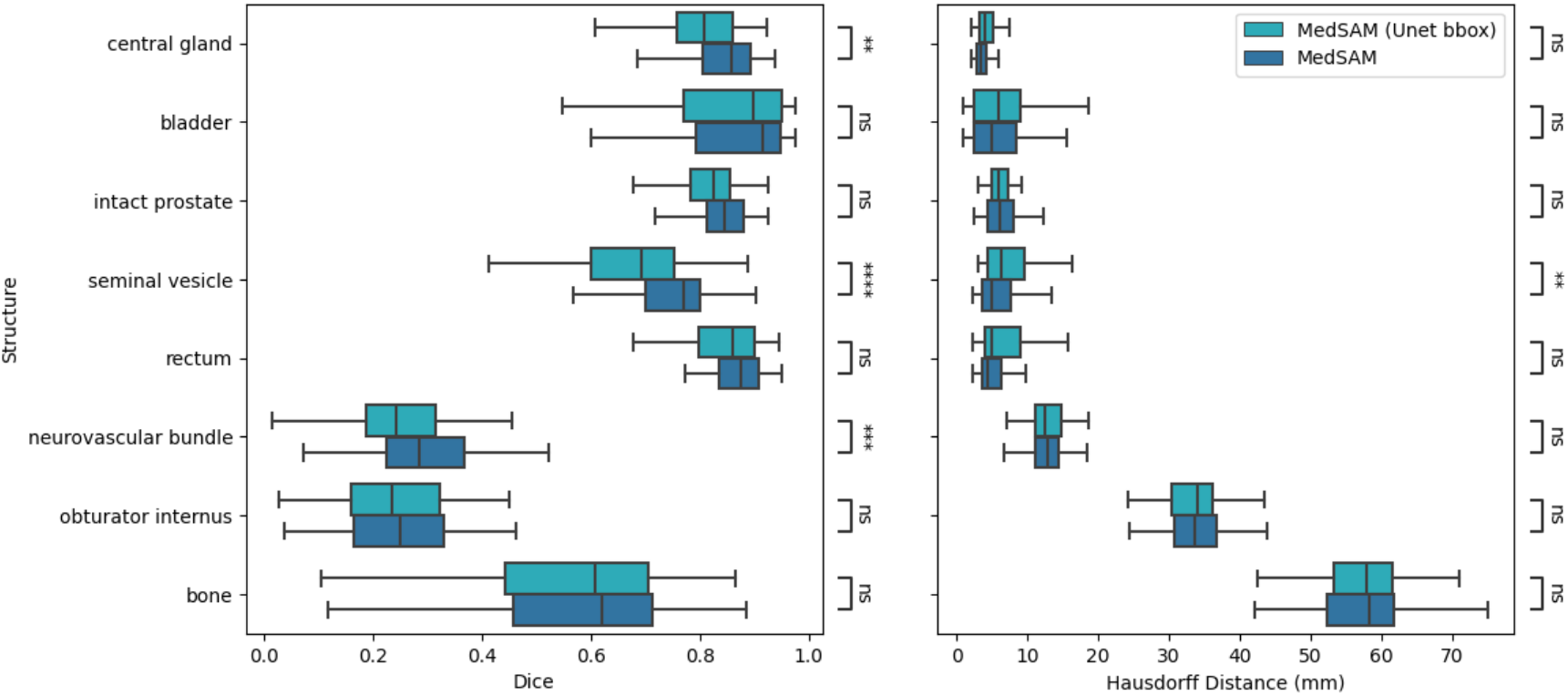
Comparison of MedSAM sensitivity to bounding box prompts derived by nnU-Net or ground truth labels. [*, **, ***, ****, ns] corresponds to statistical significance of α = [0.05, 0.01, 0.001, 0.0001, No Significance]

As shown in Table 1 columns 5 and 6, isometric expansion of both the ground truth derived bounding boxes and the nnU-Net derived bounding boxes by 5 pixels showed a very minor increase in mean Dice score for all structures but were only found to be statistically significant in the case of central gland, neurovascular bundle, and obturator internus. No significant differences in Hausdorff distances were observed.

### 3.3 Impact of specialized refinement

After being fine-tuned using a subset of our total Pelvic MR dataset, the lite MedSAM model can perform on par with the benchmark nnU-Net, as shown in *Figure 4*. Whereas out-of-the-box lite MedSAM struggled with disjoint object segmentation, the fine-tuned model has successfully learned to segment these structures more effectively. While most structures showed comparable performance between the refined lite MedSAM model and the benchmark nnU-Net in either mean Dice score or mean Hausdorff distance, LiteMedSAM showed advantage in segmenting the seminal vesicle better. *Figure 5* shows the improvement of LiteMedSAM after fine-tuning.

**Figure 4:**
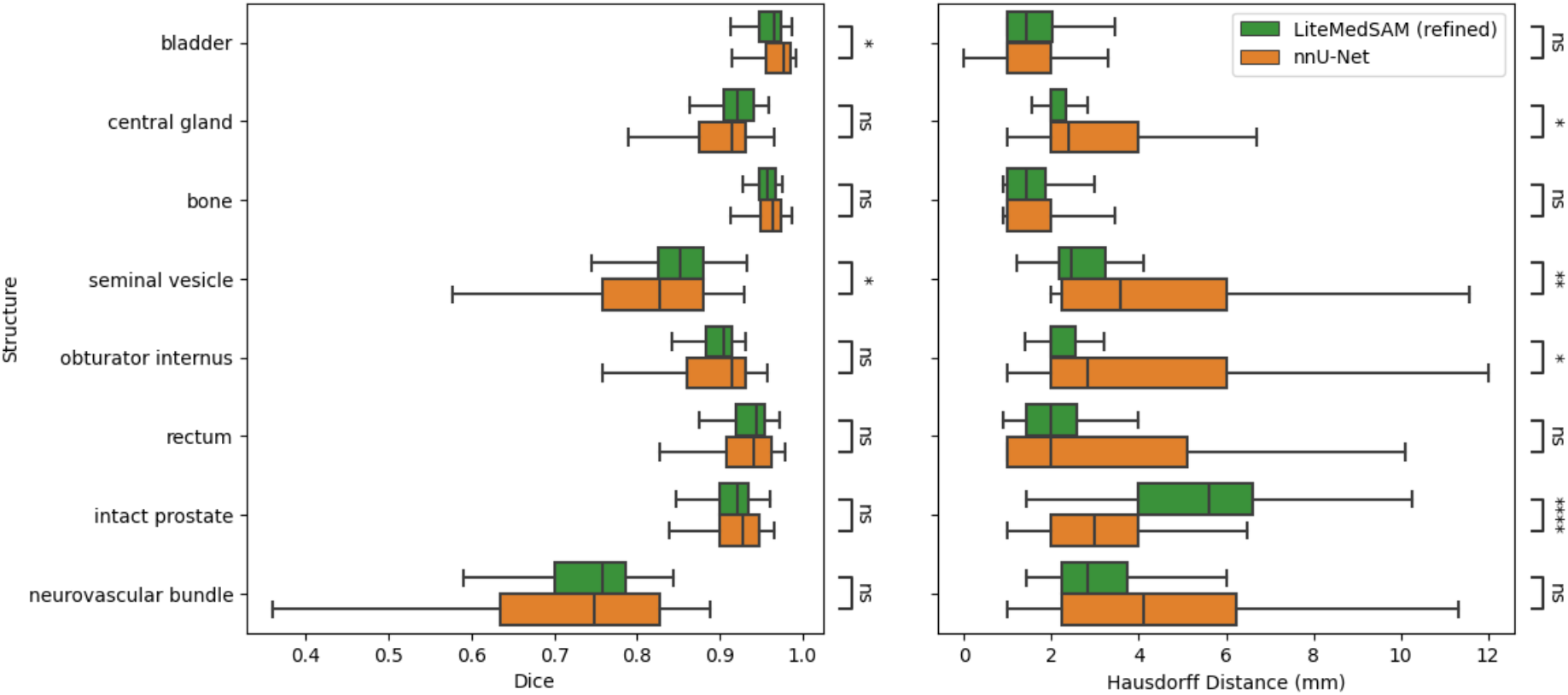
Comparison of Dice scores and Hausdorff distances between refined LiteMedSAM and nnU-Net. [*, **, ***, ****, ns] corresponds to statistical significance of α = [0.05, 0.01, 0.001, 0.0001, No Significance].

**Figure 5:**
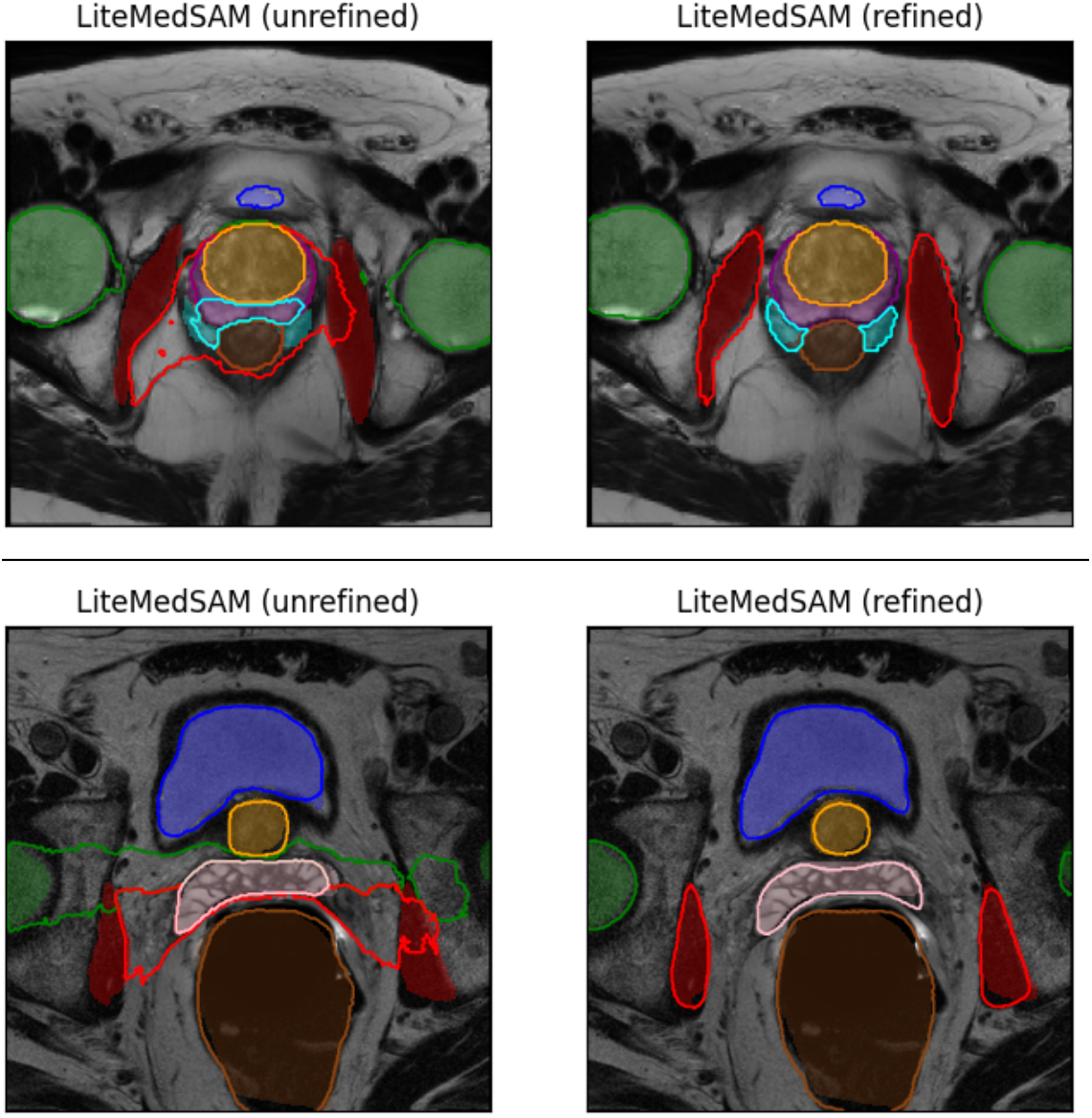
Visual comparison of LiteMedSAM before and after fine-tuning for the same two patients as shown in Figure 2.

## 4. Discussion and conclusions

In this study, we evaluated the performance of MedSAM and its light version LiteMedSAM, for the segmentation of anatomical structures in pelvic MR images. We investigated the influence of different bounding box prompts on MedSAM segmentations. More importantly, upon observing the general inferior performance on the specific task, we performed specialized fine-tuning and assessed its effectiveness. Our results indicate that out-of-the-box MedSAM and LiteMedSAM exhibit suboptimal performance compared to state-of-the-art models. We observed that that the MedSAM models have a particularly difficult time segmenting objects that are non-convex or non-elliptical despite possessing relatively well-defined boundaries. This is likely because MedSAM was originally trained on a variety of modalities which included pathology images and dermoscopy, and a variety of segmentation tasks, including cellular and molecular, which may bias the segmentation decoder to contiguous convex shapes. We also found that MedSAM is challenged by segmenting multiple, disjointed objects.

Fine-tuning lite MedSAM with a subset of our dataset yielded promising results, with the fine-tuned model performing comparably with nnU-Net, our benchmark model, especially for disjoint object segmentation. This suggests that MedSAM’s foundation model can adapt effectively to specific task and anatomy with targeted training.

While our preliminary investigation shows that varying prompting using the current bounding box input has low impact on the segmentation result, it is expected that a more flexible and enabling scheme, such as wider shape atlas or customized masking^2^, could provide implicit guidance to the underlying task and could offer higher performance gain. In addition, the current work performed specialized refinement on the decoder portion of MedSAM to achieve comparable performance to the benchmark nnU-Net. It is expected that more sophisticated refinement, such as introducing modifiers to the deep layers of the encoder portion of MedSAM may further improve performance.

Overall, our study highlights the necessity and importance of specialized fine tuning to make large foundation models like MedSAM and LiteMedSAM to be clinically relevant and useful.

